# Cheminformatics analysis of multi-target structure-activity landscape of environmental chemicals binding to human endocrine receptors

**DOI:** 10.1101/2023.10.06.561266

**Authors:** Shanmuga Priya Baskaran, Ajaya Kumar Sahoo, Nikhil Chivukula, Kishan Kumar, Areejit Samal

## Abstract

In human exposome, environmental chemicals can target, and disrupt different endocrine axes, ultimately leading to several endocrine disorders. Such chemicals, termed endocrine disrupting chemicals (EDCs), can promiscuously bind to different endocrine receptors and lead to varying biological endpoints. Thus, understanding the complexity in molecule-receptor binding of environmental chemicals can aid in the development of robust toxicity predictors. Towards this, the ToxCast project has generated the largest resource on the chemical-receptor activity data for environmental chemicals that were screened across various endocrine receptors. However, the heterogeneity in the multi-target structure-activity landscape of such chemicals is not yet explored. In this study, we systematically curated the chemicals targeting 8 human endocrine receptors, their activity values and biological endpoints from the ToxCast chemical library. We employed dual-activity difference and triple-activity difference maps to identify single-, dual-, and triple-target cliffs across different target combinations. We annotated the identified activity cliffs through matched molecular pair (MMP) based approach, and observed that a small fraction of activity cliffs form MMPs. Further, we structurally classified the activity cliffs and observed that R-group cliffs form the highest fraction among the cliffs identified in various target combinations. Finally, we leveraged the mechanism of action (MOA) annotations to analyze structure-mechanism relationships, identified strong MOA-cliffs and weak MOA-cliffs, for each of the 8 endocrine receptors. Overall, insights from this first study analyzing the structure-activity landscape of environmental chemicals targeting multiple human endocrine receptors, will likely contribute towards the development of better toxicity prediction models for characterizing the human chemical exposome.

## Introduction

The human exposome encompasses exposure to all environmental factors, and understanding the adverse effects of such exposures on human health is a key goal of environmental science in the 21^st^ century.^1^ In particular, certain environmental chemicals have been observed to interact and disrupt the normal functioning of human endocrine system, and termed as endocrine disrupting chemicals (EDCs).^2–4^ EDCs can target several endocrine axes or organs, where their selective binding to different endocrine receptors leads to different adverse outcomes such as metabolic, reproductive, neurological disorders, and cancers.^2–6^ Therefore, analyzing the multimodal nature of EDC-receptor binding can enable us to link the various adverse effects caused by the EDCs.

In pharmacology, the concept of promiscuity in molecule binding has aided in a better understanding of drug targets, and in the design of novel polypharmacological drugs.^7,8^ A similar understanding of the complexity in molecule-receptor binding of toxic chemicals^9^ can aid in the development of robust toxicity predictors. In this direction, the ToxCast project^10^ has screened nearly 10,000 environmental chemicals across multiple human receptors, and generated several chemical-receptor activity datasets.^11,12^ However, the heterogeneity in the structure-activity landscape of chemicals targeting multiple receptors have not been explored in the ToxCast chemical space.

The methodology to analyze the heterogeneity in the structure-activity landscape of chemicals has been extensively developed for a single-target chemical space^13–17^, but similar efforts are limited for multi-target chemical space. In particular, Bajorath and colleagues have primarily used network-like representations to analyze the promiscuous chemicals in the context of drug discovery research.^18–23^ Independently, Medina-Franco and colleagues had extended the concept of structure-activity similarity (SAS) map to identify multi-target activity cliffs in the drug-relevant chemical space.^24^ They proposed activity-difference maps (dual-activity difference (DAD) and triple-activity difference (TAD) maps) to analyze and identify the single-, dual- and triple-target cliffs present in the chemical space.^25,26^ Importantly, these activity-difference maps aid in the comparison in the direction of the structure-activity relationship (SAR) among chemicals forming multi-target activity cliffs. However, such methodology has not been used to analyze the multi-target toxic chemical space.

In this study, we leveraged the activity-difference map based approach to analyze the structure-activity landscape of chemicals targeting several human endocrine receptors. To achieve this, we systematically retrieved the chemical activity values and the corresponding biological endpoints across 8 human endocrine receptors from ToxCast. We employed both the DAD and TAD maps to analyze and identify the single-, dual- and triple-target cliffs across chemicals targeting all combinations of receptors. Further, we used the matched molecular pair (MMP) based approach to annotate the identified activity cliffs. Subsequently, we leveraged the structural information to classify the chemical pairs forming activity cliffs. We also analyzed the heterogeneity in the structure-mechanism relationships of chemicals targeting the different receptors, and identified mechanism of action (MOA) cliffs. In sum, the present study is the first attempt in analyzing the heterogeneity in the structure-activity landscape of toxic chemicals targeting multiple human endocrine receptors.

## Results and discussion

### Exploration of structure-activity landscape of chemicals targeting multiple human endocrine receptors

Our main objective is to analyze the heterogeneity in the structure-activity landscape of chemicals targeting multiple human endocrine receptors. To achieve this, we systematically obtained the agonist and antagonist chemical data from ToxCast project for 8 human endocrine receptors (Table S1), namely androgen receptor (AR), estrogen receptor alpha (ERα), estrogen receptor beta (ERβ), glucocorticoid receptor (GR), peroxisome proliferator-activated receptor delta (PPARδ), peroxisome proliferator-activated receptor gamma (PPARγ), progesterone receptor (PR), and thyroid stimulating hormone receptor (TSHR) (Methods; Table S2-S3). We further generated 28 dual-target combinations (Table S4-S5) and 56 triple-target combinations (Table S6-S7) in each of the agonist and antagonist datasets (Methods). We considered structurally similar chemicals in each of these combinations for further analysis (Methods).

Among the 28 dual-target agonist datasets, we observed that the activity values (pAC_50_) of chemicals targeting both GR and PPARγ showed the highest correlation (Pearson coefficient 0.75), while those targeting TSHR and ERβ showed the lowest correlation (Pearson coefficient -0.16) (Table S4). This suggests that the structure-activity landscape of chemicals targeting GR and PPARγ are more similar compared to other pairs of targets. Similarly, among the antagonist datasets, we observed activity values of chemicals targeting PPARδ and PPARγ showed highest correlation (Pearson coefficient 0.8), while those targeting PPARδ and ERβ showed the lowest correlation (Pearson coefficient -0.04) (Table S5), suggesting chemicals targeting PPARδ and PPARγ have similar structure-activity landscapes.

### Identification of single-, dual- and triple-target activity cliffs among chemicals in agonist dataset

We employed triple-activity difference (TAD) map and dual-activity difference (DAD) map based approach to identify single-, dual-, and triple-target cliffs among the generated 56 triple-target and 28 dual-target combinations in agonist dataset (Methods). The DAD map approach aids in the identification of chemical pairs forming activity cliffs against two targets (dual-target activity cliffs), whereas the TAD map approach, which is a combination of three DAD maps, additionally aids in the identification of chemical pairs forming activity cliffs against three targets (triple-target cliffs).^26^ For example, chemicals targeting AR, ERα, and ERβ showed the highest fraction of triple-target activity cliffs (13 triple-target cliffs among 200 chemical pairs; Figure 1a; Table S6). Notably, all the 13 triple-target cliffs identified in AR-ERα-ERβ TAD map show a common trend of inverse SAR with respect to AR-ERα and AR-ERβ targets, and similar SAR with ERα-ERβ targets. These 13 triple-target cliffs are formed by 14 chemicals, and ClassyFire^27^ categorized all of these chemicals under the Superclass “Lipids and lipid-like molecules”. Figure 1b shows triple-target cliffs formed by chemical pairs [CAS:4245-41-4, CAS:434-22-0] and [CAS:68-22-4, CAS:965-90-2], where the direction of SAR with respect to AR receptor is inverse with that of ERα and ERβ receptors.

**Figure 1:**
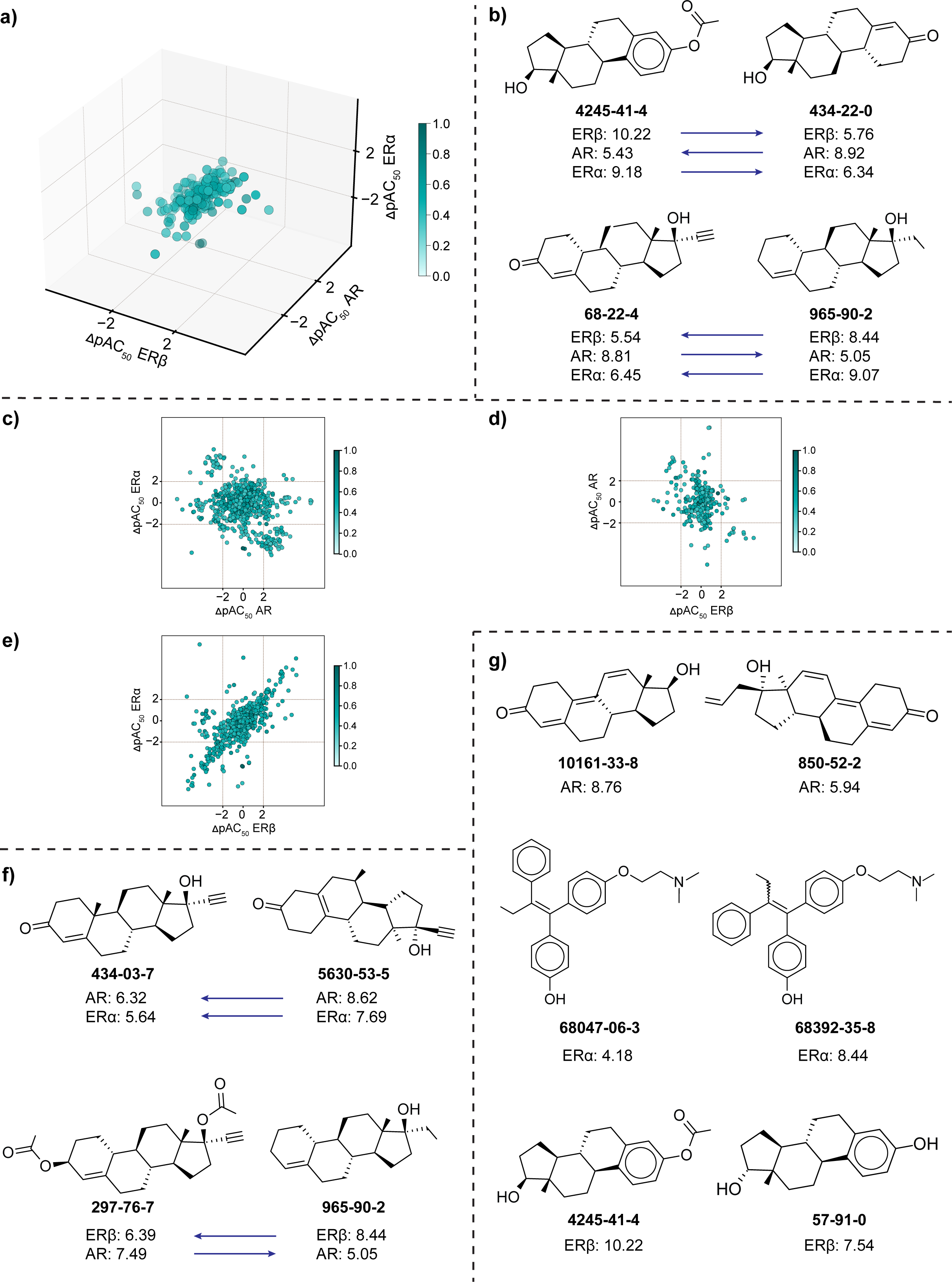
Analysis of structure-activity landscape of chemicals in the agonist dataset. **(a)** TAD map for common and structurally similar chemicals targeting AR, ERα and ERβ receptors. Each axis of the TAD map represents the values of activity difference between the chemicals in each pair for one receptor. The color gradient represents the similarity between chemicals in each pair (where the darker shade represents higher structural similarity). **(b)** Chemical pairs forming triple-target cliffs, where the arrow denotes the direction of SAR. **(c-e)** DAD maps corresponding to each pair of receptors considered in the TAD map. The axis represents the receptors, and the color gradient represents the similarity between chemicals in each pair (where the darker shade represents higher structural similarity). **(f)** Chemical pairs forming dual-target cliffs, where the arrow denotes the direction of SAR. **(g)** Chemical pairs forming single target cliffs with respect to each of the receptors considered in the TAD map.

We further split the AR-ERα-ERβ TAD map into three DAD maps, namely AR-ERα (Figure 1c), AR-ERβ (Figure 1d) and ERα-ERβ (Figure 1e), to identify the dual-target and single-target cliffs. Among the three DAD maps, the AR-ERα DAD Map showed the highest fraction of dual-target and single-target cliffs (91 dual-target cliffs and 174 single-target cliffs among 859 chemical pairs; Table S4). Among the 91 dual-target cliffs identified in AR-ERα DAD map, 10 have similar SAR and the remaining have inverse SAR. Notably, these 91 dual-target cliffs are formed by 51 chemicals, among which 46 chemicals are classified under the Superclass “Lipids and lipid-like molecules”. Figure 1f shows an example of a similar SAR dual-target cliff formed by chemical pair [CAS:434-03-7, CAS:5630-53-5] with respect to AR and ERα receptors, and an inverse SAR dual-target cliff formed by chemical pair [CAS:297-76-7, CAS:965-90-2] with respect to AR and ERβ receptors. Figure 1g shows examples of single-target cliffs with respect to each of AR [CAS:10161-33-8, CAS:850-52-2], ERα [CAS:68047-06-3, CAS:68392-35-8], and ERβ [CAS:4245-41-4, CAS:57-91-0] receptors.

### Identification of single-, dual- and triple-target activity cliffs among chemicals in antagonist dataset

Similar to the agonist dataset, we employed TAD and DAD map based approach to identify the single-, dual-, and triple-target cliffs among the structurally similar chemicals from the 56 triple-target and 28 dual-target combinations in antagonist dataset (Methods). Chemicals targeting AR, PPARδ, and PPARγ showed the highest fraction of triple-target activity cliffs (66 triple-target cliffs among 1060 chemical pairs; Figure 2a; Table S7). Notably, 62 of 66 triple-target cliffs identified in AR-PPARδ-PPARγ TAD map show a common trend of similar SAR with respect to AR-PPARδ, AR-PPARγ and PPARδ-PPARγ targets. The 66 triple-target cliffs are formed by 60 chemicals, among which 43 are classified under the Superclass “Benzenoids”. Figure 2b shows triple-target cliffs formed by chemical pairs [CAS:603-33-8, CAS:76-87-9] with similar SAR with respect to all the 3 receptors (AR, PPARδ, and PPARγ) and [CAS:22978-25-2, CAS:4822-44-0], where the direction of SAR with respect to PPARγ receptor is inverse with that of AR and PPARδ receptors.

**Figure 2:**
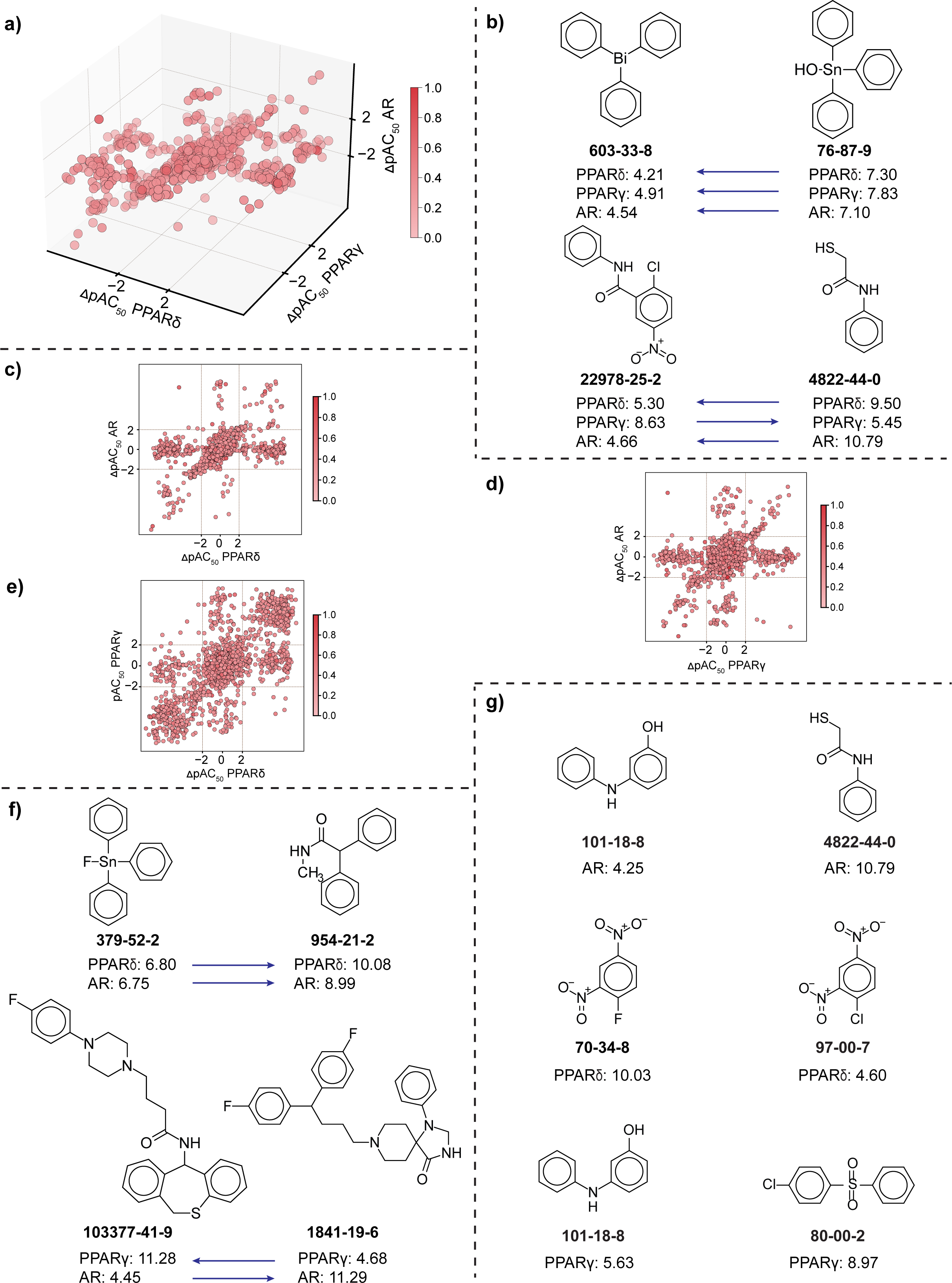
Analysis of structure-activity landscape of chemicals in the antagonist dataset. **(a)** TAD map for common and structurally similar chemicals targeting AR, PPARδ, and PPARγ receptors. Each axis of the TAD map represents the values of activity difference between the chemicals in each pair for one receptor. The color gradient represents the similarity between chemicals in each pair (where the darker shade represents higher structural similarity). **(b)** Chemical pairs forming triple-target cliffs, where the arrow denotes the direction of SAR. **(c-e)** DAD maps corresponding to each pair of receptors considered in the TAD map. The axis represents the receptors, and the color gradient represents the similarity between chemicals in each pair (where the darker shade represents higher structural similarity). **(f)** Chemical pairs forming dual-target cliffs, where the arrow denotes the direction of SAR. **(g)** Chemical pairs forming single target cliffs with respect to each of the receptors considered in the TAD map.

Further, upon splitting the AR-PPARδ-PPARγ TAD map into three corresponding DAD maps (Figure 2c-e), we observed that PPARδ-PPARγ DAD map showed the highest fraction of dual-target cliffs (621 out of 2226 pairs) while AR-PPARδ showed the highest fraction of single-target cliffs (353 out of 1521 pairs) (Table S5). Among the 621 dual-target cliffs from PPARδ-PPARγ DAD map (Figure 2e), 608 show similar SAR and 13 show inverse SAR. These 621 dual-target cliffs are formed by 342 chemicals, among which 198 chemicals are classified under the Superclass “Benzenoids”. The high percentage of similar SAR among PPARδ-PPARγ dual-target cliffs can be attributed to the high correlation between activity values of the chemicals targeting these two receptors. Figure 2f shows an example of a similar SAR dual-target cliff formed by chemical pair [CAS:379-52-2, CAS:954-21-2] with respect to AR and PPARδ receptors, and an inverse SAR dual-target cliff formed by chemical pair [CAS:103377-41-9, CAS:1841-19-6] with respect to AR and PPARγ receptors. Figure 2g shows examples of single-target cliffs with respect to each of AR [CAS:101-18-8, CAS:4822-44-0], PPARδ [CAS:70-34-8, CAS:97-00-7], and PPARγ [CAS:101-18-8, CAS:80-00-2] receptors.

### Matched molecular pair (MMP) based annotation of the identified activity cliffs

We leveraged the matched molecular pair (MMP) based approach to annotate the activity cliffs identified in both the agonist and antagonist datasets via DAD and TAD maps (Methods; Table S8). Among the different receptor combinations in the agonist dataset, the dual-target combination of ERα-ERβ receptors (Figure 3a) show the highest fraction of MMPs while the triple-target combination of GR-PPARγ-TSHR receptors (Figure 4a) show the highest fraction of MMPs. Similarly, among the different receptor combinations in the antagonist dataset, the dual-target combination of ERβ-PPARγ receptors (Figure 3b) show the highest fraction of MMPs while the triple-target combination of ERα-GR-TSHR receptors (Figure 4b) show the highest fraction of MMPs. Figure 3c displays the examples of MMP formed by the dual-target combination of PPARδ-PR in agonist dataset (CAS:705-60-2, CAS:102-96-5) and PPARδ-PR in antagonist dataset (CAS:52918-63-5, CAS:39515-40-7).

**Figure 3:**
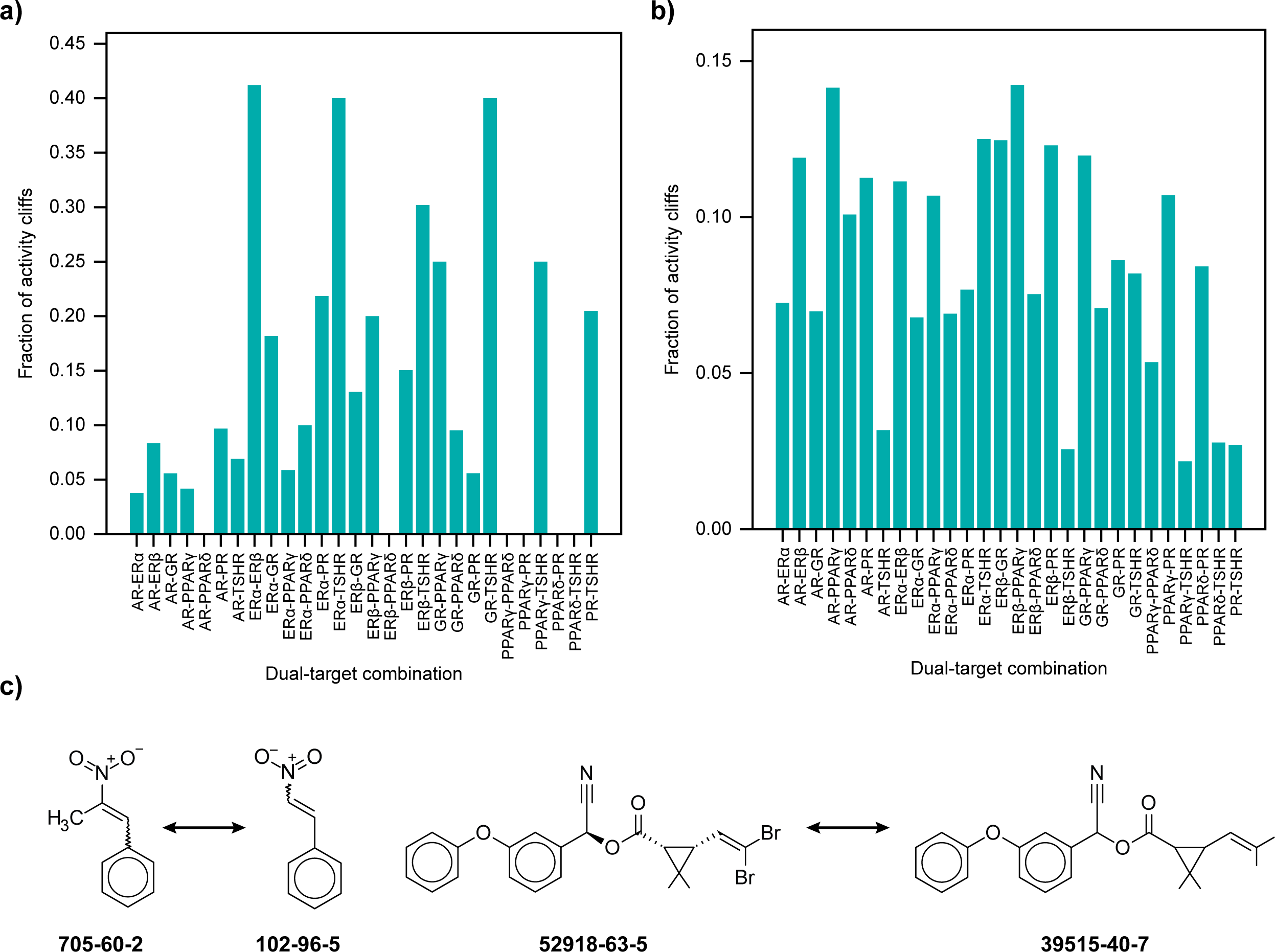
Exploration of MMPs among the activity cliffs identified via DAD maps. **(a)** Distribution of MMPs among activity cliffs identified in the agonist dataset via DAD maps. 23 **(b)** Distribution of MMPs among activity cliffs identified in the antagonist dataset via DAD maps. **(c)** Chemical pairs forming MMPs in agonist and antagonist datasets.

**Figure 4:**
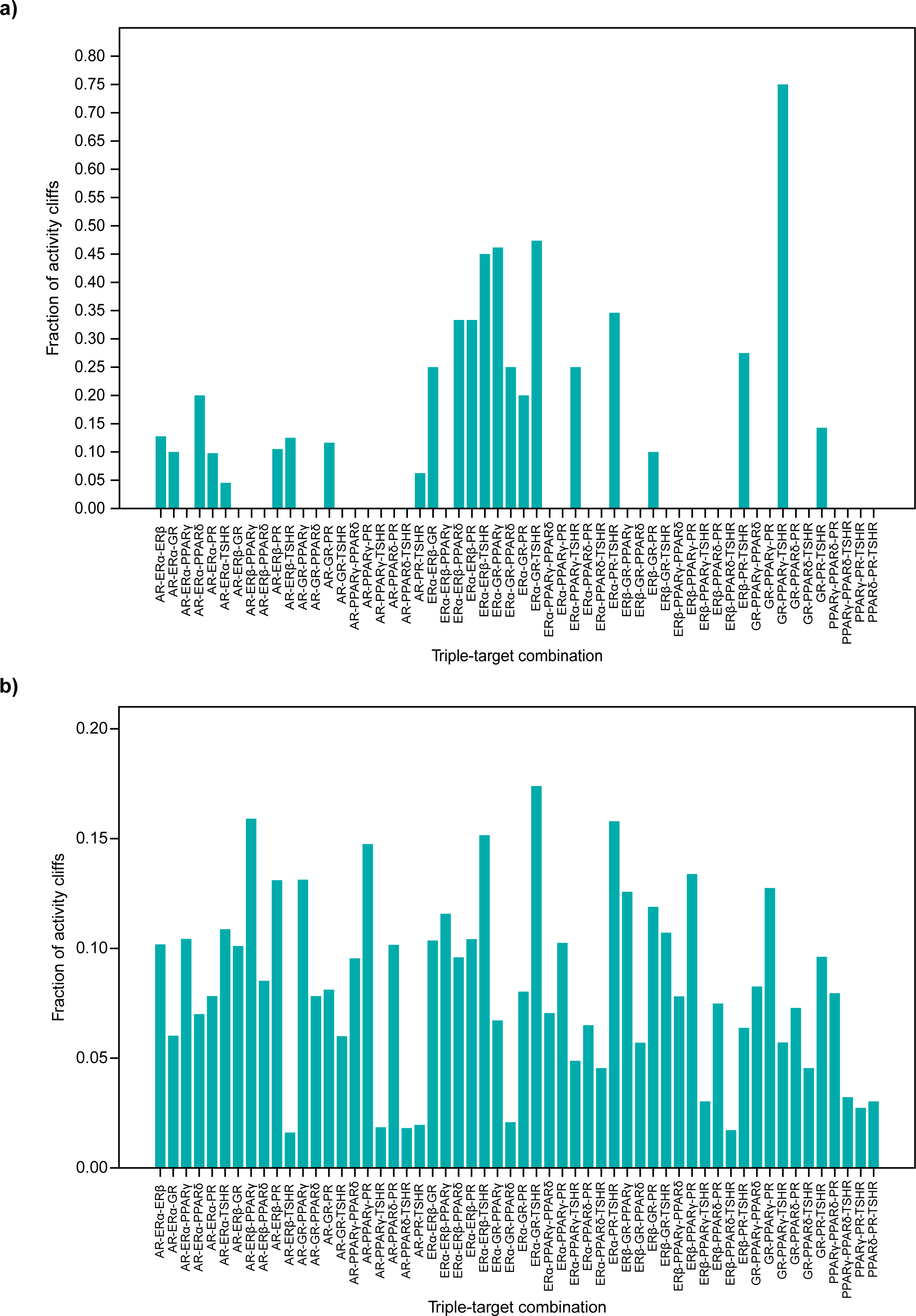
Exploration of MMPs among the activity cliffs identified via TAD maps. **(a)** Distribution of MMPs among activity cliffs identified in the agonist dataset via TAD maps. **(b)** Distribution of MMPs among activity cliffs identified in the antagonist dataset via TAD maps.

### Structural classification of the identified activity cliffs

By leveraging the structural features of the chemicals, we independently classified the activity cliffs identified in agonist and antagonist datasets via DAD and TAD maps (Methods, Table S9). We observed that in most of the dual-target or triple-target combinations, the fraction of the R-group cliffs is highest. Among the different receptor combinations in the agonist dataset, the dual-target combination of AR-PR and ERβ-PR showed all six structural classifications (Figure 5a), while the triple-target combination of ERα-ERβ-PR showed a maximum of 5 structural classifications (Figure 6a). Similarly, among the different combinations in the antagonist dataset, the dual-target combinations of AR-PPARγ, ERα-PPARγ, and PPARγ-PR (Figure 5b), and the triple-target combinations of AR-ERα-PPARγ, AR-PPARγ-PR, and ERα-PPARγ-PR (Figure 6b) showed all 6 structural classifications. Figure 5c shows examples of chemical pairs forming 6 structural classifications, namely chirality cliff (formed by CAS:28434-00-6 and CAS:584-79-2 across 13 DAD maps in antagonist dataset), topology cliff (formed by CAS:452-86-8 and CAS:95-71-6 across 15 DAD maps in antagonist dataset), R-group cliff (formed by CAS:637-03-6 and CAS:705-60-2 across 18 DAD maps in antagonist dataset), scaffold cliff (formed by CAS:16320-04-0 and CAS:6533-00-2 across 4 DAD maps in agonist dataset and 3 DAD maps in antagonist dataset), scaffold/topology cliff (formed by CAS:434-03-7 and CAS:5630-53-5 across 5 DAD maps in agonist dataset), and scaffold/R-group cliff (formed by CAS:4584-57-0 and CAS:493-52-7 across 15 DAD maps in antagonist dataset).

**Figure 5:**
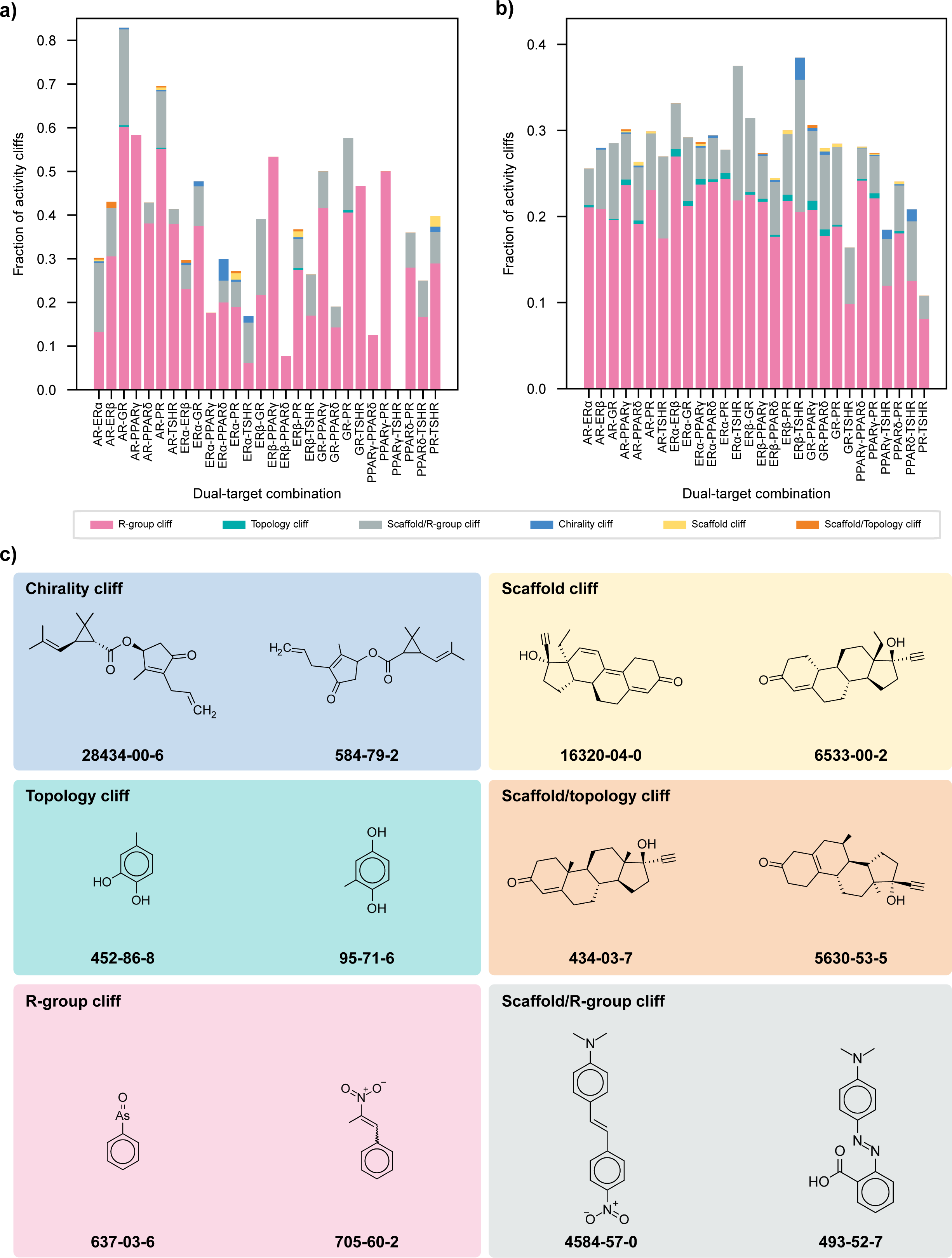
Classification of activity cliffs identified via DAD maps. **(a)** Distribution of different structural classifications among activity cliffs identified in agonist dataset via DAD maps. **(b)** Distribution of different structural classifications among activity cliffs identified in antagonist dataset via DAD maps. **(c)** Activity cliff pairs forming different structural classes.

**Figure 6:**
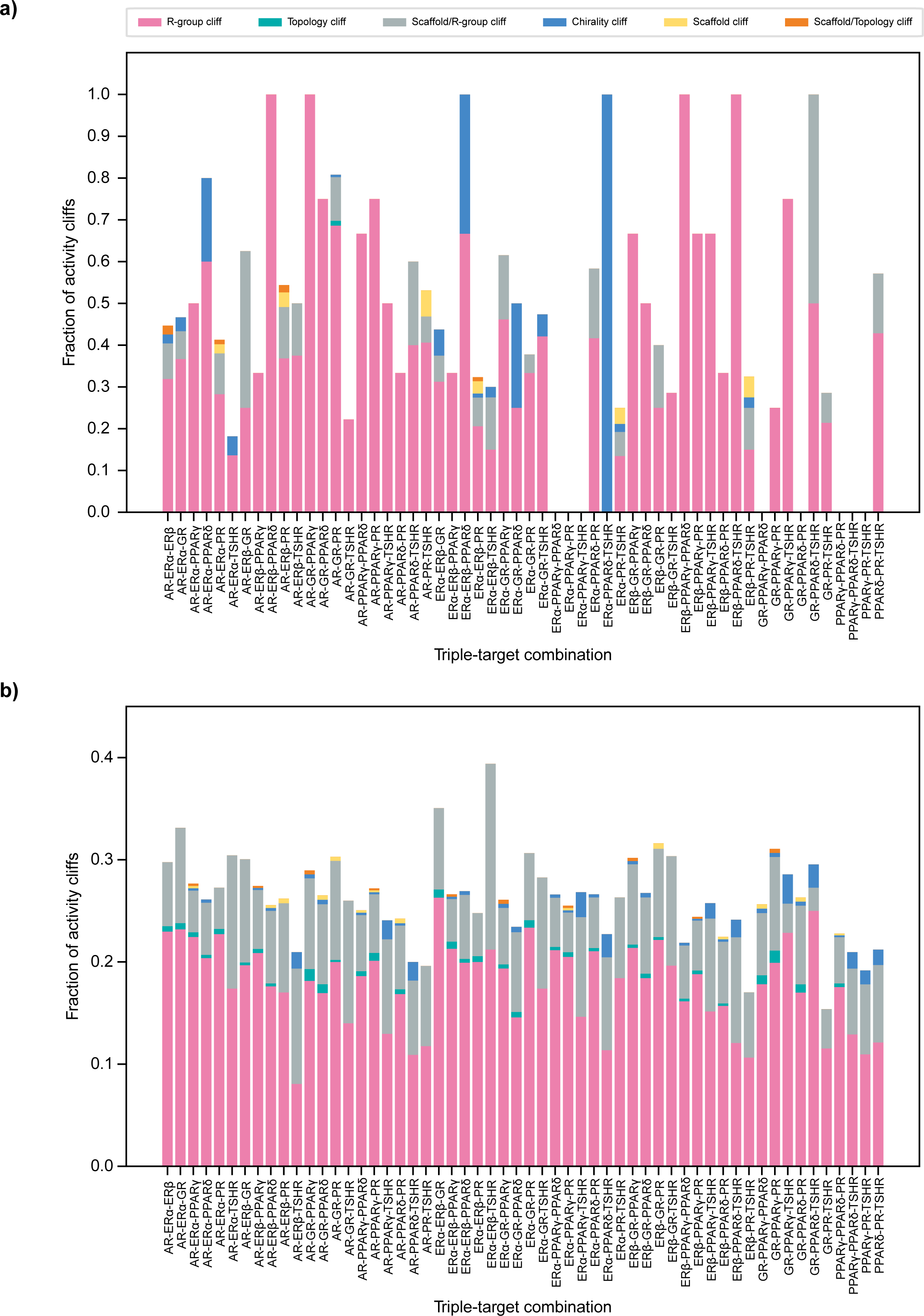
Classification of activity cliffs identified via TAD maps. **(a)** Distribution of different structural classifications among activity cliffs identified in agonist dataset via TAD maps. **(b)** Distribution of different structural classifications among activity cliffs identified in antagonist dataset via TAD maps.

### Identification of strong and weak mechanism of action (MOA) cliffs

In addition to analyzing the structure-activity landscape, we analyzed the heterogeneity in the structure-mechanism relationship of chemicals with respect to each of the 8 human endocrine receptors. We shortlisted the common chemicals between agonist and antagonist dataset for each receptor, and thereafter, identified strong and weak mechanism of action (MOA) cliffs using their MOA annotations (Methods; Table S10). Figure 7a shows the distribution of the strong MOA-cliffs, weak MOA-cliffs, and same MOA across each of the 8 receptors. Among the 8 receptors, PPARγ has the highest fraction of strong MOA-cliffs (8 of 83 MOA pairs) and weak MOA-cliffs (42 of 83 MOA pairs), while PR has the highest number of strong MOA-cliffs (54 MOA pairs) and weak MOA-cliffs (807 MOA pairs). Figure 7b displays examples of strong MOA-cliff (formed by CAS:10540-29-1 and CAS:68047-06-3), weak MOA-cliff (formed by CAS:68047-06-3 and CAS:68392-35-8), and same MOA (formed by CAS:102-96-5 and CAS:5153-67-3) with respect to PPARγ receptor.

**Figure 7:**
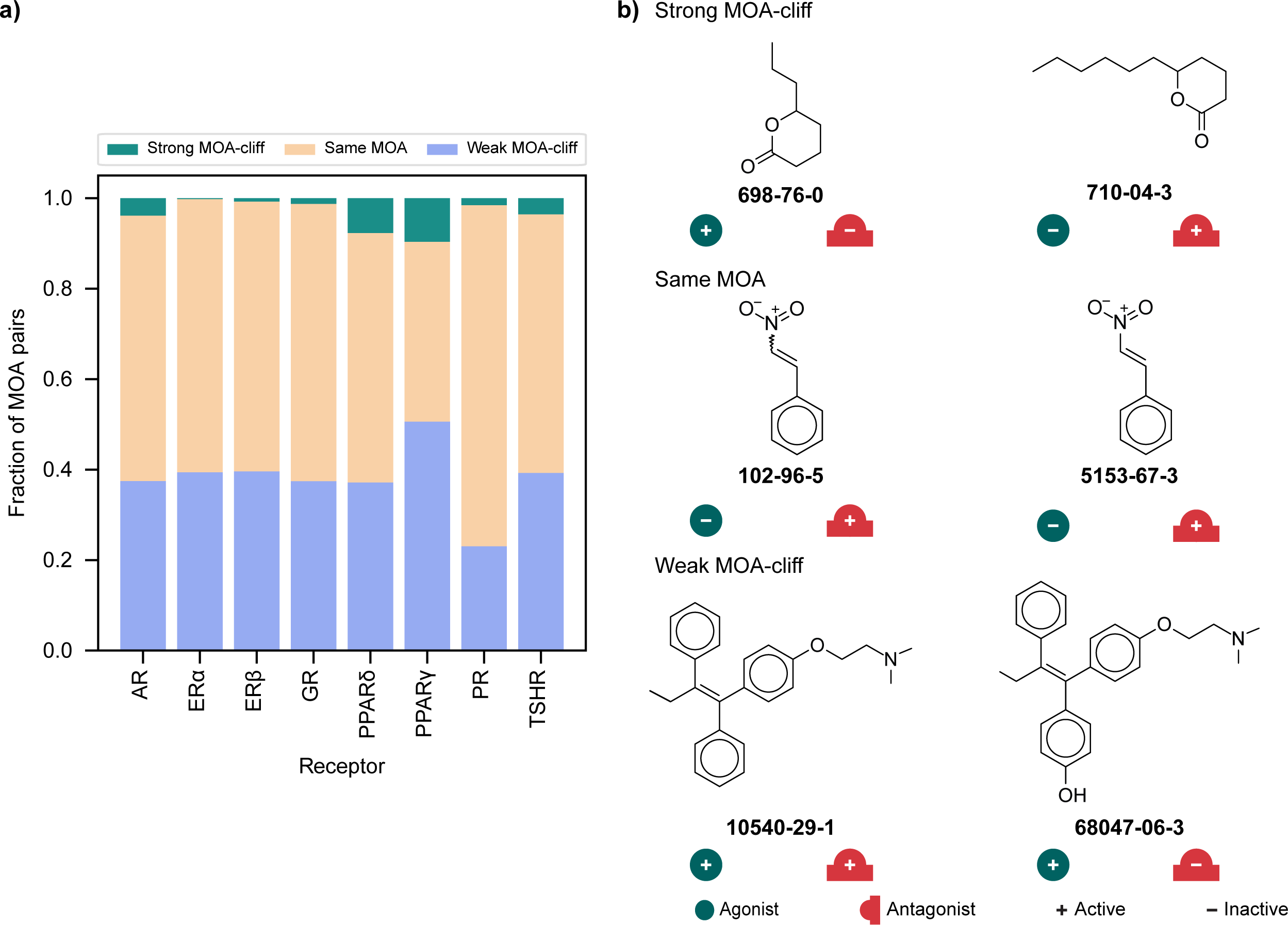
Identification of various mechanism of action (MOA) cliffs. **(a)** Distribution of strong MOA-cliffs, weak MOA-cliffs and same MOA across chemicals targeting the 8 human endocrine receptors. **(b)** Chemical pairs forming strong MOA-cliff, same MOA, and weak MOA-cliff.

### Conclusions

In this study, we systematically analyzed the structure-activity landscape of environmental chemicals targeting multiple human endocrine receptors. First, we curated the list of chemicals binding to 8 human endocrine receptors from the ToxCast chemical library, and obtained their corresponding activity values and biological endpoints. Then, we leveraged different cheminformatics tools such as dual-activity difference (DAD), and triple-activity difference (TAD) maps, and identified chemical pairs forming single-, dual-, and triple-target cliffs. Further, we structurally annotated the activity cliffs using the matched molecular pair (MMP) based approach, and observed that a low fraction (0.08) of the activity cliffs form MMPs. We also structurally categorized the activity cliff pairs and observed that the R-group cliff category has the highest fraction among activity cliffs of various target combinations. Finally, we leveraged the mechanism of action (MOA) annotations and explored the structure-mechanism relationships among chemicals targeting each of the 8 receptors. To the best of our knowledge, this is the first study that explores and analyzes the structure-activity landscape of environmental chemicals targeting multiple human endocrine receptors.

ToxCast chemical library is the largest chemical resource that has screened various environmental chemicals across different cell lines, and quantitatively cataloged the corresponding biological interactions. Among the curated list of 3829 chemicals from ToxCast analyzed in this study, 312 chemicals have been cataloged as EDCs in DEDuCT^5^, and 474 chemicals are documented as high production volume (HPV) chemicals by OECD HPV^28^ and US HPV.^29^ We also observed a low overlap of chemicals binding to all 8 human endocrine receptors in each of the agonist (20 chemicals) and antagonist (100 chemicals) datasets. This suggests that the extent of promiscuity in the binding of environmental chemicals (assessed by ToxCast project) to these 8 endocrine receptors is low.

However, the present study does not address the molecular mechanisms underlying the formation of various multi-target activity cliffs. Further, we are restricted to only 8 human endocrine receptors, due to lack of high confidence datasets. Nonetheless, this study highlights the presence of multi-target activity cliffs among environmental chemicals targeting different human receptors. Overall, we expect the findings from this study will aid in the development of well-informed machine learning based toxicity prediction models, and contribute towards human chemical exposome research.

## Methods

### Curated dataset of chemicals targeting multiple endocrine receptors

In this study, our main objective is to analyze the activity landscape of chemicals that can target multiple endocrine receptors in humans. To this end, we leveraged the chemical dataset from the high-throughput Tox21 assays (assay source identifier 7) with level 5 and 6 preprocessing within ToxCast version 3.5.^30^ Tox21 captures 146 assay endpoints spread across various cell lines and targets. To ensure a high confidence chemical dataset specific to human endocrine receptors, we filtered the Tox21 data to obtain assay endpoints that: (i) are primary readouts; (ii) are performed on human cell lines; (iii) have corresponding agonist or antagonist endpoint annotations; (iv) are not follow-up assays; (v) independently target a single human endocrine receptor. Based on these criteria, we identified assays corresponding to 16 endpoints across 8 human endocrine receptors (Table S1).

Thereafter, we used an in-house R script to obtain the chemical information from ToxCast for these 16 assay endpoints. In particular, we filtered chemicals that were annotated as representative chemicals (gsid_rep = 1) and have a reported activity value (modl_ga is present). Next, we accessed the two-dimensional (2D) structures of chemicals from ToxCast version 3.5 or PubChem.^31^ Further, we used MayaChemTools^32^ to remove the salts, invalid molecules, mixtures and duplicated chemicals. Moreover, based on our previous studies^16,17^, we computed the Bemis-Murcko scaffolds^33^ of chemicals and removed the linear molecules. Finally, we curated and compiled 8 human endocrine receptor specific agonist datasets (Table S2) and antagonist datasets (Table S3) containing the CAS or PubChem identifiers, activity values, and endpoint annotations (active or inactive), which were further considered for various analyses.

### Computation of chemical similarity and activity difference

We computed the structural similarity between any pair of chemicals based on Tanimoto coefficient of their corresponding ECFP4 (Extended connectivity fingerprints with diameter 4) chemical fingerprints. The activity difference between any pair of chemicals is given by the difference in their pAC_50_ values, where pAC_50_ is the negative logarithm of AC_50_ value in molar concentration. The obtained chemical datasets from ToxCast consist of chemical activities mentioned in terms of modl_ga values, which is the logarithm of AC_50_ value in micromolar concentration. We used the following formulae to obtain the corresponding pAC_50_ value of chemicals:

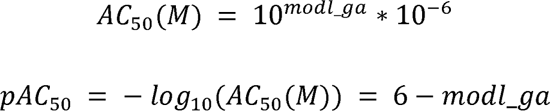

Finally, we computed the activity difference between two chemicals against a particular target T using the following formula:

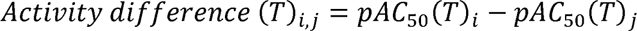

### Identification of activity cliffs based on dual activity-difference (DAD) maps and triple activity-difference (TAD) maps

We independently analyzed the structure-activity relationship (SAR) of chemicals in agonist and antagonist datasets using the dual activity-difference (DAD) maps and triple activity-difference (TAD) maps.^25,26^ DAD map is a 2D representation where the axes denote the activity difference between chemicals against two different targets, and each point on the plot denotes a chemical pair (Figure 8). We obtained all possible combinations of targets from each dataset, and considered the common chemicals in these combinations (Table S4-5). Further, in each of these combinations, we obtained structurally similar chemical pairs as those having a similarity value greater than or equal to three standard deviations from median of the similarity distribution. For each of these chemical pairs, we computed their activity difference by preserving their sign, and plotted them on a DAD map. To identify significant differences in activity values, we set an activity difference threshold of -2 and 2 on each axis, and divided the DAD map into 5 zones, namely zone I to zone V (Figure 8). The chemical pairs in zone I show a similar trend of differences in activity ([ΔpAC_50_(T1) and ΔpAC_50_(T2)] > 2 or < -2) against both targets, denoting similar SAR. The chemical pairs in zone II show an inverse trend of differences in activity (ΔpAC_50_(T1) > 2 and ΔpAC_50_(T2) < -2, or vice versa) against the two targets, denoting inverse SAR. Additionally, chemical pairs in zone I and zone II are referred to as dual-target cliffs.^26^ The chemical pairs in zone III and zone IV show significant difference in activity against only one of the two targets, and are referred to as single-target cliffs.^26^ The chemical pairs in zone V show no significant difference in activity against either of the targets, and hence do not form activity cliffs.

**Figure 8:**
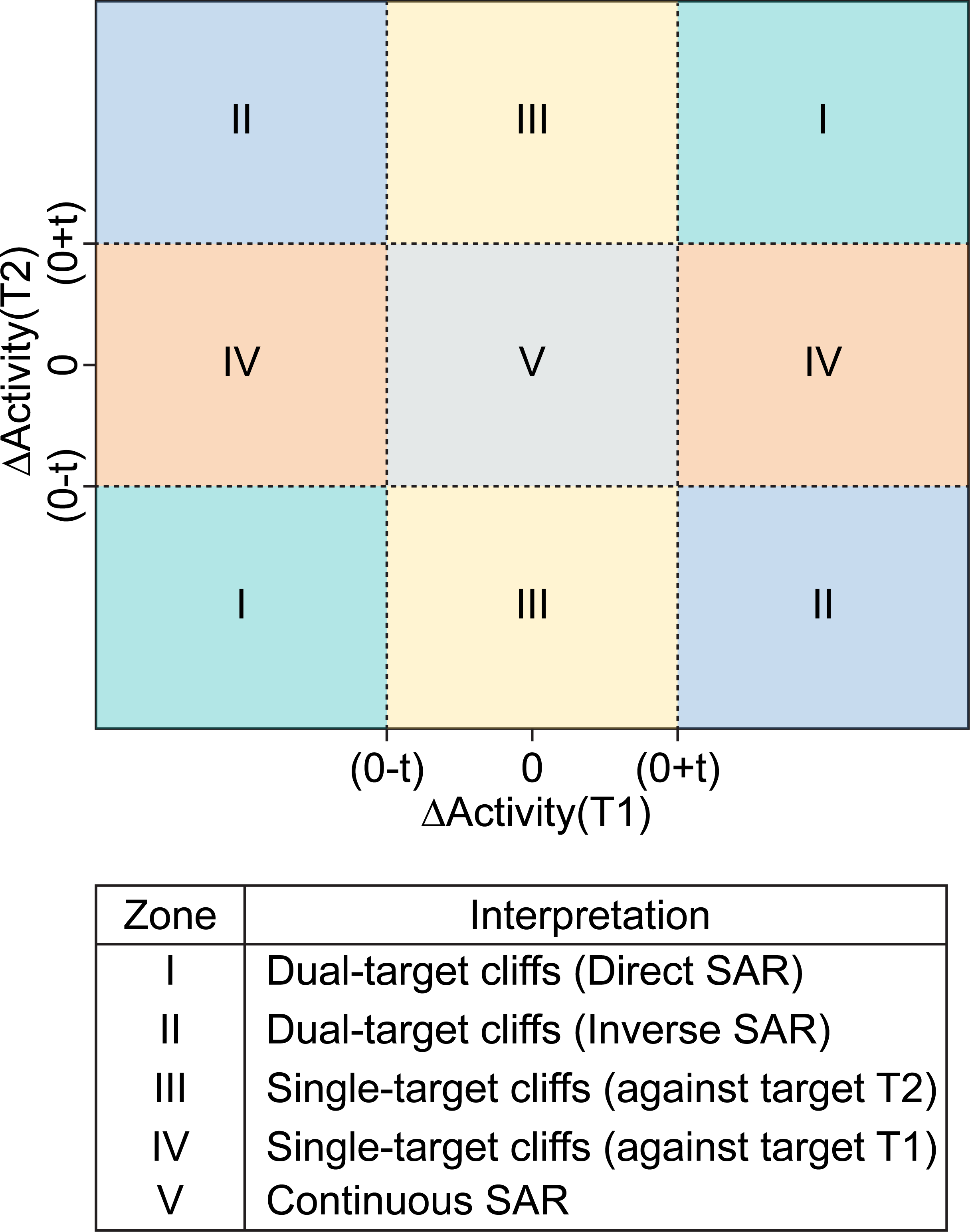
Prototype of the dual-activity difference (DAD) map against targets T1 and T2 where ‘t’ is the activity difference threshold. The DAD map can be divided into 5 zones (I to V) and the interpretation of the 5 zones is tabulated below the DAD map.

TAD map is a 3D representation where the axes denote the activity difference between chemicals against three different targets, and each point on the plot denotes a chemical pair (Figure 9). Similar to the DAD map approach, we obtained structurally similar chemical pairs for all possible combinations of three targets, and computed their corresponding differences in activity values. To identify significant differences in activity values, we set an activity difference threshold of +2 and -2 along each axis, and identified single-target, dual-target and triple-target cliffs (Figure 9).

**Figure 9:**
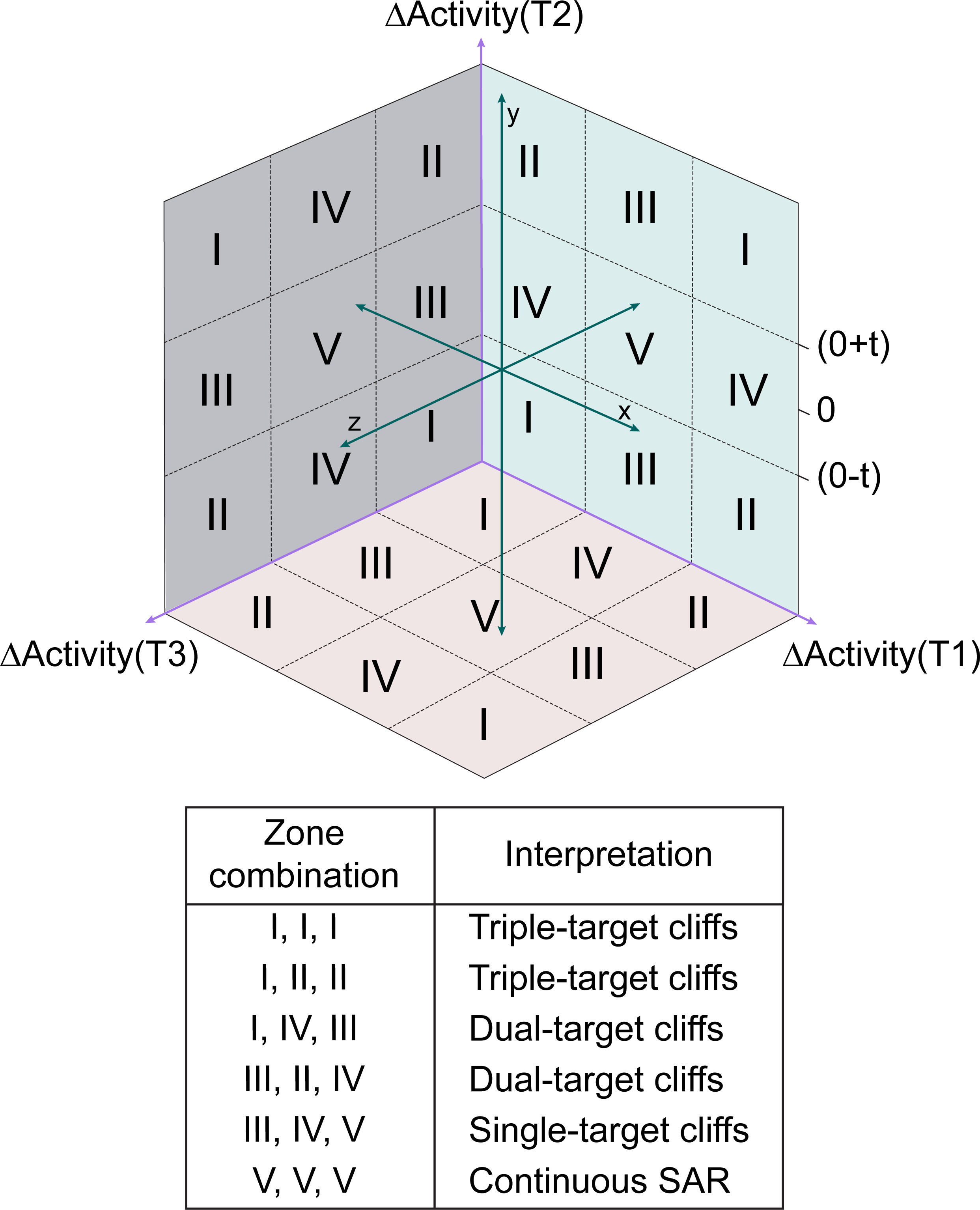
Prototype of the triple-activity difference (TAD) map against targets T1, T2, and T3, where ‘t’ is the activity difference threshold. The interpretation of the various zone combinations of the TAD map is tabulated below the TAD map.

### Annotation of activity cliffs based on matched molecular pairs (MMPs)

Matched Molecular Pairs (MMPs) are chemical pairs that structurally differ at a single site. It is a substructure-based approach that is descriptor-independent, metric-free and chemically intuitive.^34^ Therefore, we employed the MMP-based approach^35^ to annotate the activity cliffs identified from DAD and TAD maps. Based on our previous work^17^, we employed the mmpdb platform^36^ to generate MMPs for each of the 16 datasets. First, we performed the fragmentation of the chemicals using mmpdb fragmentation module with ‘none’ value for both maximum number of non-hydrogen atoms and maximum number of rotatable bonds arguments. Then, we generated an exhaustive list of MMPs by using the mmpdb index module with ‘none’ value for maximum number of non-hydrogen atoms in the variable fragment argument. Further, we obtained the size restricted MMPs using the following criteria^17^:

i. The difference in number of heavy atoms of the exchanged fragments in the transformation should be ≤ 8.
ii. The constant part of a MMP should be at least twice the size of each fragment in the transformation.
iii. The number of heavy atoms (non-hydrogen atoms) of each fragment in the transformation should be ≤ 13.
iv. If a chemical pair has multiple MMPs, the transformation which has the least heavy atom difference in the exchanged fragments is considered.

### Structural classification of activity cliffs

We provided structural classification of activity cliffs identified from DAD and TAD maps by considering information on molecular scaffolds, R-groups, R-group topology, and chirality of chemical structures.^15^ Based on our previous work^17^, we developed a python workflow that employs RDKit^37^ to classify the activity cliffs into the following seven classes:

i. Chirality cliff: activity cliff pairs whose scaffolds, R-groups, and R-group topology are same.
ii. Topology cliff: activity cliff pairs whose R-group topologies are different, while their scaffolds and R-groups are same.
iii. R-group cliff: activity cliff pairs whose R-groups are different, while their scaffolds are same.
iv. Scaffold cliff: activity cliff pairs whose scaffolds are different, while their cyclic skeletons, R-groups, and R-group topologies are same.
v. Scaffold/topology cliff: activity cliff pairs whose scaffolds and R-group topologies are different while their cyclic skeletons and R-groups are same.
vi. Scaffold/R-group cliff: activity cliff pairs whose scaffolds and R-groups are different, while their cyclic skeletons are same.
vii. Unclassified: activity cliff pairs whose both scaffolds and cyclic skeletons are different.

### Analysis of the structure-mechanism relationships of endocrine receptor binding chemicals

Apart from the heterogeneity in their structure-activity landscape, chemicals can show heterogeneity in their structure-mechanism relationships leading to mechanism of action (MOA) cliffs.^17,34^ Based on our previous work^17^, we first identified the common chemicals between the agonist and antagonist datasets for each of the 8 human receptors. Then, for each receptor, we filtered out the chemicals that were inactive in both the agonist and antagonist assay endpoints (hit_c value is 0) and shortlisted structurally similar chemical pairs whose Tanimoto coefficient was greater than or equal to 3 standard deviations from the median of the similarity distribution. Finally, based on the MOA annotations, we classified the chemical pairs (MOA pairs) into three categories:

i. Strong MOA-cliff: chemical pairs for which the MOA annotations are opposite in both agonist and antagonist assay endpoint.
ii. Same MOA: chemical pairs for which MOA annotations are same in both agonist and antagonist assay endpoint.
iii. Weak MOA-cliff: chemical pairs which couldn’t be categorized as either Strong MOA-cliff or Same MOA.

## Author Contributions

**Shanmuga Priya Baskaran:** Conceptualization, Data Compilation, Data Curation, Formal Analysis, Software, Visualization, Writing; **Ajaya Kumar Sahoo:** Conceptualization, Data Compilation, Data Curation, Formal Analysis, Software, Visualization, Writing; **Nikhil Chivukula:** Formal Analysis, Writing; **Kishan Kumar:** Formal Analysis, Visualization; **Areejit Samal:** Conceptualization, Supervision, Formal Analysis, Writing.

## Acknowledgements

The authors thank Dhiraj Kumar for stimulating discussions. Areejit Samal acknowledges funding from the Department of Atomic Energy (DAE), Government of India [Apex project to The Institute of Mathematical Sciences (IMSc), Chennai] and the Max Planck Society, Germany [Max Planck Partner Group in Mathematical Biology]. The funders have no role in the study design, data collection, data analysis, manuscript preparation, or decision to publish.

## Supporting Information

**Table S1:** Curated list of 8 human endocrine receptors considered in this study. For each receptor, the table provides the agonist assay endpoint identifier, agonist assay endpoint name, number of chemicals considered for the agonist assay, antagonist assay endpoint identifier, antagonist assay endpoint name and number of chemicals considered for the antagonist assay, from the ToxCast chemical library.

**Table S2:** Curated list of chemicals in the agonist dataset for the 8 receptors from ToxCast chemical library. For each chemical, the table provides the chemical identifier, Canonical SMILES computed using Open Babel, computed pAC_50_ value against the 8 receptors. Note, if the activity value for a chemical against a given receptor is not provided by ToxCast, then the corresponding cell is left empty. Values are shown up to 2 decimal places.

**Table S3:** Curated list of chemicals in the antagonist dataset for the 8 receptors from ToxCast chemical library. For each chemical, the table provides the chemical identifier, Canonical SMILES computed using Open Babel, computed pAC_50_ value against the 8 receptors. Note, if the activity value for a chemical against a given receptor is not provided by ToxCast, then the corresponding cell is left empty. Values are shown up to 2 decimal places.

**Table S4:** List of 28 DAD maps formed by dual-target combinations (shown by the receptor name, separated by ’-’) among 8 receptors in the agonist dataset. For each DAD map, the table provides number of chemical pairs formed by common structurally similar chemicals, computed Pearson coefficient between activity values (pAC_50_) of the chemicals targeting the two receptors, computed Pearson coefficient between activity difference (ΔpAC_50_) for chemical pairs targeting the two receptors, number and fraction of chemical pairs forming single-target cliffs, number and fraction of chemical pairs forming dual-target cliffs, and number and fraction of total activity cliffs (combination of both single-target and dual-target cliffs). Values are shown up to 2 decimal places.

**Table S5:** List of 28 DAD maps formed by dual-target combinations (shown by the receptor name, separated by ’-’) among 8 receptors in the antagonist dataset. For each DAD map, the table provides number of chemical pairs formed by common structurally similar chemicals, computed Pearson coefficient between activity values (pAC_50_) of the chemicals targeting the two receptors, computed Pearson coefficient between activity difference (ΔpAC_50_) for chemical pairs targeting the two receptors, number and fraction of chemical pairs forming single-target cliffs, number and fraction of chemical pairs forming dual-target cliffs, and number and fraction of total activity cliffs (combination of both single-target and dual-target cliffs). Values are shown up to 2 decimal places.

**Table S6:** List of 56 TAD maps formed by triple-target combinations (shown by the receptor name, separated by ’-’) among 8 receptors in the agonist dataset. For each TAD map, the table provides number of chemical pairs formed by common structurally similar chemicals, number and fraction of chemical pairs forming single-target cliffs, number and fraction of chemical pairs forming dual-target cliffs, number and fraction of chemical pairs forming triple-target cliffs, and number and fraction of total activity cliffs (combination of single-target, dual-target, and triple-target cliffs). Values are shown up to 2 decimal places.

**Table S7:** List of 56 TAD maps formed by triple-target combinations (shown by the receptor name, separated by ’-’) among 8 receptors in the antagonist dataset. For each TAD map, the table provides number of chemical pairs formed by common structurally similar chemicals, number and fraction of chemical pairs forming single-target cliffs, number and fraction of chemical pairs forming dual-target cliffs, number and fraction of chemical pairs forming triple-target cliffs, and number and fraction of total activity cliffs (combination of single-target, dual-target, and triple-target cliffs). Values are shown up to 2 decimal places.

**Table S8:** List of matched molecular pairs (MMPs) identified among the chemicals in agonist or antagonist datasets analyzed in this study. For each MMP, the table provides the chemical identifiers, chemical SMILES (after performing canonicalization in RDKit and removing stereoisomer information, which was used then as input SMILES in mmpdb platform in order to generate MMPs), generated constant part and transformation between the chemical structures. Further, the table provides the list of dual-target and triple-target combinations (separated by ’|’) for a chemical pair forming an activity cliff.

**Table S9:** List of activity cliffs identified among the chemicals in agonist or antagonist datasets analyzed in this study. For each activity cliff, the table provides the chemical identifiers, structural classification, and list of dual-target and triple-target combinations (separated by ’|’) where the activity cliff is identified.

**Table S10:** Classification of chemicals targeting each of the 8 receptors based on their mechanism of action (MOA) annotation reported in the corresponding agonist and antagonist assay endpoint. For each receptor, the table provides number of common chemicals which have at least one active endpoint in either agonist or antagonist assay, computed chemical similarity threshold by using the Tanimoto coefficient between the ECFP4 fingerprints, number of strong MOA-cliff, weak MOA-cliff, and same MOA. Values are shown up to 2 decimal

